## Supplemental Materials for "Uncertainty-driven regulation of learning and exploration in adolescents: A computational account"

### Supplementary Material

#### Supplementary Text 1. Deviations from preregistration

*Regression analyses (estimation task)*. Consistent with our preregistration, we tested the effects of trial, noise level and age group on certainty and learning rate. However, we deviated from our preregistration in *how* we modeled the effects of trial. According to our preregistration, we would fit power functions to each participant's timecourse of certainty ratings and learning rates—separately for the two noise conditions—and then perform regression analyses on the estimated power-function parameters. After preregistering, however, we realized that this power-function approach was too specific for our research questions, and that a more flexible and parsimonious approach would be to model the linear and quadratic effects of trial (which also allowed the use of multilevel models).

*Mediation analysis (estimation task)*. We preregistered a related but different analysis, namely a regression of learning rate on certainty and age group. However, as both certainty and learning rate were strongly related to trial number, we reasoned that it would be more informative to control for trial in this analysis. Therefore, we used a multilevel moderated mediation analysis that simultaneously tests (i) whether trial-to-trial changes in certainty are predictive of learning rate when controlling for trial number (path *b* of the mediation model); (ii) whether certainty mediates the relationship between trial number and learning rate; and (iii) whether each of these effects differ between age groups.

*Exclusion of outlier learning rates in the behavioural analyses of learning rate (estimation task).* We did not preregister the exclusion of trials on which the estimated learning rate exceeded the 99th percentile or was lower than the 1st percentile. These most extreme learning-rate estimates likely resulted from occasional typing errors (e.g., when a participant accidentally typed '3' instead of '83')—as verified by inspection of the estimation data—something we did not anticipate when preregistering. Note that this exclusion criterion is usual in studies where learning rates are directly estimated from single-trial estimation data (Vaghi et al., 2017).

*Analyses of learning rate as a function of age within the adolescent group.* We did not preregister the analysis reported in the “Learning rate decreases over the course of early adolescence” section; hence this analysis should be considered exploratory.

*Computational models.* We preregistered the standard reinforcement learning and Kalman filter models, but not the asymmetric reinforcement learning and reinforcement learning/Pearce-Hall hybrid models. We added these latter two models to examine potential alternative explanations for the observed age-related differences in learning rate and choice behaviour, as suggested by the reviewers.

*Analyses of the choice task.* The reported analyses of the choice data were not preregistered, because we initially intended to report the results from this task separately, focusing on a different question (see http://aspredicted.org/blind.php?x=av4td4). However, after preregistering, we realized that the estimation and choice tasks yield complementary information about uncertainty-driven changes in reinforcement learning; hence could provide converging evidence regarding developmental changes. Therefore, we decided to analyze the choice data in a similar way as the estimation data, and report both sets of results together.

**Supplementary Text 2. Optimal adaptation of learning rate in noisy but static environments**

When outcomes are drawn from a static Gaussian distribution—as in our experimental tasks—the best estimate of the mean outcome after *t* observations (i.e., on trial

$t$+1) is simply the average of all outcomes, $O$, that have been observed so far (from trial 1 through trial $t$):

$$E_{t+1} = (O_1 + O_2 + \cdots + O_t)/t = \frac{1}{t}\sum_{i=1}^{t} O_i \qquad \text{[S1]}$$

This 'sample-average' method can be achieved using a reinforcement-learning algorithm in which the learning rate, $\alpha$, is reduced over trials according to $\alpha_t = \frac{1}{t}$, as illustrated by the dotted line in Fig 2B in the main text. This can be seen by rewriting equation S1 as follows:

$$E_{t+1} = \frac{1}{t}\sum_{i=1}^{t} O_i = \frac{1}{t}\left(O_t + \sum_{i=1}^{t-1} O_i\right) = \frac{1}{t}\left(O_t + (t-1)E_t\right) = \frac{1}{t}\left(O_t + tE_t - E_t\right)$$

$$= \frac{O_t}{t} + E_t - \frac{E_t}{t} = E_t + \frac{1}{t}(O_t - E_t)$$

This corresponds to the delta-rule learning algorithm from equation 1.2 in the main text, with $\frac{1}{t}$ is trial-specific $\alpha$, and $O_t - E_t$ is prediction error $\delta_t$.

Note that this optimal learning-rate regime is approximated by the Kalman filter when (i) the initial prior variance, relative to the noise variance, is set at a very high value (such that the initial learning rate approaches 1) and (ii) the drift variance, relative to the noise variance, is 0 (such that the asymptotic learning rate approaches 0).

**Supplementary Text 3: Model recovery analysis**

*Procedure.* We simulated data on each task with each of our models. Each simulated dataset consisted of a group of 25 synthetic participants. The number of blocks and trials per synthetic participant were the same as in the real datasets. For the estimation task we only simulated data from the low-noise condition of the task (standard deviation of outcome-generating distribution = 4)

For each simulation, the hyperparameters governing the model's group-level distributions—from which the individual-level parameters were drawn—were sampled randomly from uniform distributions. We matched the range of the uniform distributions to the range of values obtained from our fits to the real data, using the minimum and maximum values of the hyperparameters' posterior medians for the two age groups and (for fits to the estimation data) noise conditions (Supplementary Table 2).

To examine to what extent the data-generating models could be recovered, we applied all models to each simulated dataset and determined the best fitting model for each dataset, using the same model-fitting and comparison procedure as used for the real data. We repeated this procedure 50 times for the estimation data, and 23-29 times for the choice data (the number of repetitions per data-generating model is indicated in parentheses in Supplementary Fig 3B). We used less repetitions for the choice-task simulations for practical reasons: the larger number of models and longer-lasting fitting procedure for this task.

*Results.* We summarize the results in confusion and inversion matrices [1]. Confusion matrices represent the probability that data simulated with a given model is best fit by each of the models, i.e., *p*(fit model | simulated model). For the estimation data, the probability that a dataset simulated with a given model was best fit by that same model, as opposed to one of the other three models, ranged from 0.94 to 1 (Supplementary Fig 3A, left panel). Thus, our procedure could distinguish the learning processes captured by the four different models applied to the estimation data with high accuracy. For the choice data, we found one case in which model-recovery failed: 63% of the datasets generated by the reinforcement learning/Pearce-Hall hybrid model + constant softmax was best fit by the standard RL model + constant softmax. Thus, when the degree of exploration was constant, our method did not identify the adjustment of learning rate according to the Pearce-Hall mechanism, but instead detected a constant learning rate. This is likely due to the low values of decay parameter $\bar{\eta}$ for the reinforcement learning/Pearce-Hall hybrid model + constant softmax (Supplementary Table 2). Such low values of $\bar{\eta}$ produce small decreases in learning rate over time, which could not be dissociated from a constant learning rate. In addition, data generated with the asymmetric RL model + dynamic softmax was best fit by that same model with a probability of .63, and by the standard RL model + dynamic softmax with a probability of .33, indicating moderate identifiability of this model. This may reflect that the values of $\alpha_+$ and $\alpha_-$ did not differ enough to dissociate asymmetric from symmetric expectation updating. For the other models, the simulated choice data was best fit by the data-generating model, as opposed to one of the other seven models, with probabilities ranging from .75 to .97 (Supplementary Fig 3B, left panel).

To more directly address the question of how to interpret our model-selection results—i.e., how confident can we be that our best-fitting models indeed generated our participants' data, for which the true underlying model was unknown—we also plotted the inversion matrices (right panels of Supplementary Fig 3). These matrices represent the probability that data that is best fit by a given model is generated by each of the models—*p*(simulated model | fit model)—and can be computed from the confusion matrices using Bayes rule, assuming a uniform prior on models [1]. For the estimation data, the probability that a dataset that was best fit by the Kalman filter (our best-fitting model in both age groups) was indeed generated by that model was .96, validating our model-selection results for the estimation task. For the choice data, the probability that datasets which were best fit by the asymmetric reinforcement learning model + dynamic softmax and the reinforcement learning/Pearce-Hall hybrid model + dynamic softmax (our best-fitting models for the adolescents and adults, respectively) were generated by these same models was .89 and .71, respectively. Thus, we can be rather confident about the model-selection results for the adolescents' choice data, but should interpret the best-fitting model for the adults' choice data with some caution. For datasets best fit by the latter model, there was a probability of .19 that they were generated by the standard reinforcement learning model + dynamic softmax. Therefore, we cannot say definitively whether, in the choice task, the adults decreased their learning rate over time according to a Pearce-Hall algorithm or used a constant learning rate. Importantly, the probability that datasets best fit by a dynamic softmax function were indeed generated by a dynamic, instead of a constant, softmax function (regardless of the learning model) ranged from .9 to .97, validating our conclusion that participants in both age groups decreased their degree of exploration over time.

**Supplementary Text 4: Parameter recovery analysis**

*Procedure.* We conducted parameter-recovery analyses for the best-fitting models for each task. For the estimation task, we compared the simulated vs. recovered Kalman filter parameters obtained from the model-recovery procedure described above. To further validate the difference in estimated $\overline{\sigma_\eta^2}$ between the two age groups, we also simulated 100 additional datasets with the Kalman filter, using the posterior medians of $\overline{\sigma_\eta^2}$ and $\overline{s_1^2}$ from our fits to the adolescent and adult data (for the low noise condition), for 50 simulations each, and fitted the Kalman filter model to each of these simulated datasets. We then computed 50 difference distributions, one for of each pair of recovered $\overline{\sigma_\eta^2}$ posteriors, and examined the proportion of pairs for which more than 95% of the difference distribution lay below 0 (i.e., the probability that a group difference was detected in the recovered data).

For the choice task, the range of simulated group-level mean parameters used in the model-recovery analyses was too narrow to examine correlations between simulated and recovered values. Therefore, we simulated 50 additional datasets with the winning model for each age group, using a wider range of parameter values. Specially, group-level mean parameters were randomly sampled from the following uniform distributions: $\overline{\alpha_+}$ and

$\overline{\alpha_{-}}$~U(.10, .90), $\overline{\alpha_{1}}$ ~U(.20, .95), $\bar{\eta}$~U(.05, .95), $\bar{\kappa}$~U(.20, .95), $\bar{\theta}$~U(.02, .30), $\bar{c}$~U(.50, .95). Note that we did not simulate values for $\bar{c}$ below .50 because these would produce a decreasing inverse temperature (increase in exploration), which does not correspond to our data or to normative behaviour. Group-level precision parameters were set to the center of the corresponding uniform distributions reported in Supplementary Table 2, in all simulations.

*Results.* The parameter-recovery results of the estimation task revealed a positive correlation between simulated and recovered values of $\overline{\sigma_{\eta}^{2}}$ (the drift variance parameter of the Kalman filter) within the range of the two age groups' estimated values ($r$ = .45, p = .001). However, the recovered values of $\overline{\sigma_{\eta}^{2}}$ were lower than the simulated values, and this bias was larger for higher simulated values (Supplementary Fig 4A, left plot). This suggests that relative, but not absolute, values of $\overline{\sigma_{\eta}^{2}}$ can be interpreted. Simulated and recovered values of $\overline{s_{1}^{2}}$ (the initial prior variance of the Kalman filter) were uncorrelated ($r$ = .05, $p$ = .72; Supplementary Fig 4A, right plot). This can be explained by the fact that different values of $s_{1}^{2}$ within our high range of simulated values (in the order of hundreds) lead to almost identical behavior. Specifically, all values of $s_{1}^{2}$ > 100 lead to initial learning rates that approach 1 (Equation 3.5 in the main text, note that $\sigma_{\varepsilon}^{2}$ was set to 1), and to an almost identical adjustment of the prior variance—and hence learning rate—on subsequent trials (Equation 3.2 in the main text). Recovered values of $\overline{\sigma_{\eta}^{2}}$ and $\overline{s_{1}^{2}}$ were uncorrelated ($r$ = .02, $p$ = 88), indicating that there was no trade-off between the two Kalman-filter parameters.

Our follow-up analysis that specifically addressed the difference in $\overline{\sigma_{\eta}^{2}}$ between the two age groups showed that the posterior median of $\overline{\sigma_{\eta}^{2}}$ from fits to the adults' estimation data (.00009) was well recovered, while the estimated value of $\overline{\sigma_{\eta}^{2}}$ from fits to the adolescents' data (.02) was strongly underestimated (Supplementary Fig 4B). Recovered $\overline{\sigma_{\eta}^{2}}$ was numerically higher for data simulated with the estimated values from the adolescent than the adult group in all simulations, but the recovered group difference was smaller than the simulated difference. For 70% of the simulations, more than 95% of the recovered difference distribution ($\overline{\sigma_{\eta}^2}$ adults – $\overline{\sigma_{\eta}^2}$ adolescents) lay below 0. These findings suggest that the 'real' value of $\overline{\sigma_{\eta}^2}$ for the adolescent group is likely to be higher—and the age-related difference in $\overline{\sigma_{\eta}^2}$ larger—than it seemed based on the model fits to the empirical data.

Regarding the choice-task models, simulated and recovered group-level mean parameters from the asymmetric reinforcement learning model + dynamic softmax—the winning model for the adolescents—were strongly correlated (Supplementary Fig 5A), and there was no evidence for bias, or for tradeoffs between parameters (all correlations between recovered parameters < .25, *p*'s > .08). For the reinforcement learning/Pearce-Hall hybrid model + dynamic softmax—the winning model for the adults—simulated and recovered hyperparameters were also correlated. However, $\bar{\eta}$, $\bar{\kappa}$ and $\bar{c}$ tended to be underestimated, whereas $\bar{\theta}$ tended to be overestimated for this model (Supplementary Fig 5B). In addition, $\bar{\kappa}$ (constant component of the learning rate) and $\bar{\theta}$ (inverse temperature halfway a task block) were negatively correlated ($r = -.43$, $p = .002$), reflecting the typical tradeoff between learning rate and inverse temperature [2].

**Supplementary Fig 1.** Control analyses on the dynamic-softmax parameters. We repeated the group comparisons of $\bar{\theta}$, $\bar{c}$, and inverse temperature reported in the main text, this time using the parameter estimates derived from the same model in both age groups: the reinforcement learning/Pearce-Hall hybrid model + dynamic softmax. **A**. Posterior distributions for the group-level central tendencies of the dynamic-softmax parameters per age group (left panels) and the corresponding difference distributions (right panels*)* for fits to the choice data (corresponding to Fig. 5D in the main text). **B**. Model's predicted inverse temperature per trial (corresponding to Fig 5E, right panel, in the main text).

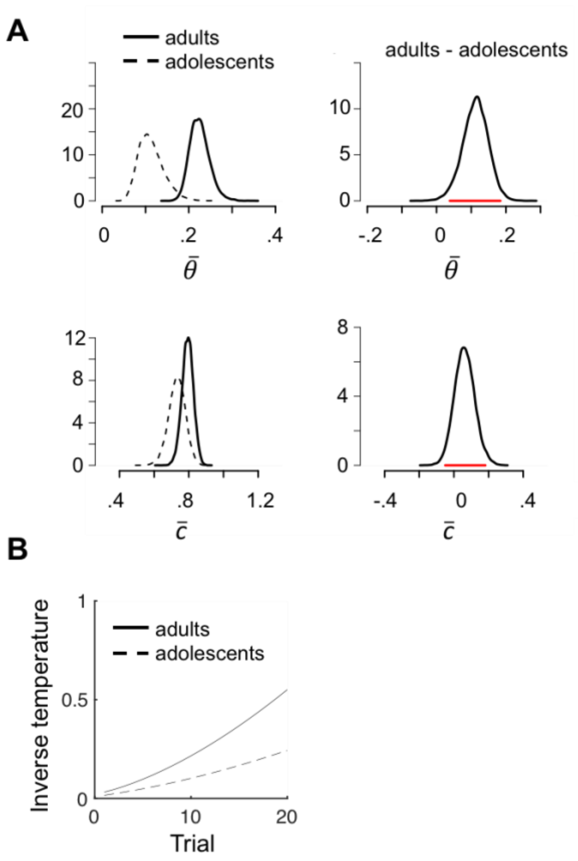

**Supplementary Fig 2**. Control analyses on the relationships between age group and model parameters $\sigma_\eta^2$, $\theta$ and $c$. We repeated the mediation analyses reported in the main text (Fig 6), this time using $\theta$ and $c$ estimates derived from the same choice-task model in both age groups **A**. Mediation models and results. **B**. Across-subject relationships between $\sigma_\eta^2$ and $\theta$, and between $\sigma_\eta^2$ and $c$ (all converted to ranks), controlled for age group (path $b$ of the mediation models). The partial correlations are $r$ = -.34, $p$ = .01 $r$ = -.47, $p$ < .001, respectively.

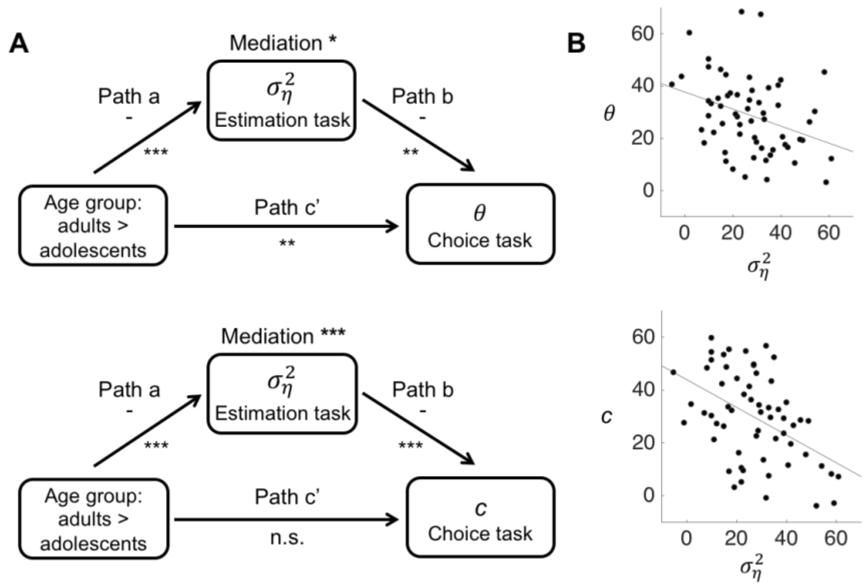

**Supplementary Fig 3.** Model-recovery results for the estimation (**A**) and choice (**B**) task. For the estimation data, we simulated 50 datasets from each model. For the choice task, the number of simulated datasets is indicated in parentheses. Models 1-4 are the reinforcement-learning, asymmetric reinforcement-learning, Kalman-filter, and reinforcement learning/Pearce-Hall hybrid models, respectively. Version a and b of each learning model (in Fig B) are combined with the constant and dynamic softmax function, respectively.

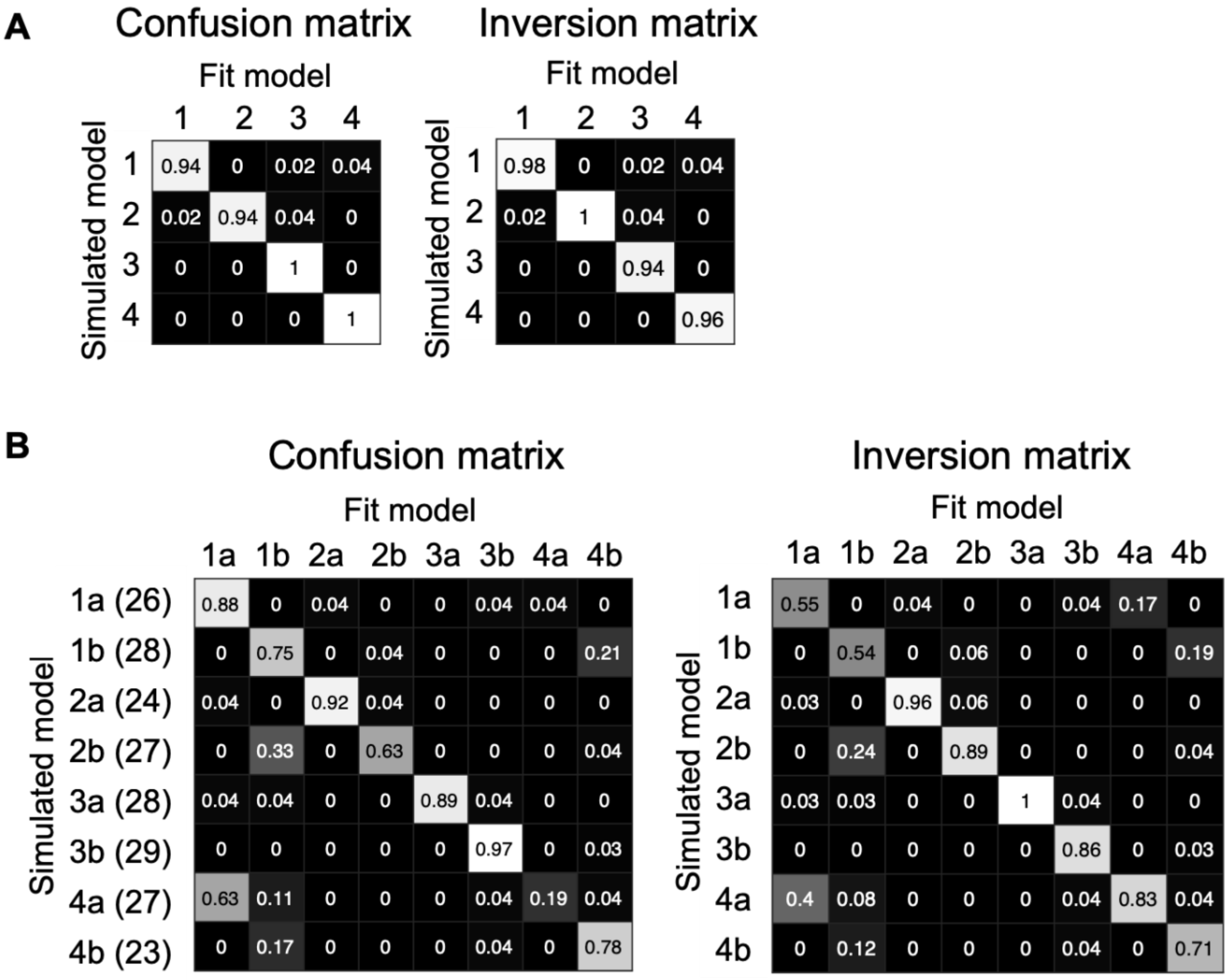

**Supplementary Fig 4**. Parameter-recovery results for the Kalman filter in the estimation task. **A**. Simulated vs. recovered $\overline{\sigma_\eta^2}$ and $\overline{s_1^2}$. Simulated values for both hyperparameters were randomly drawn from a uniform distribution which range matched the range of values from our fits to the real data. Black lines are regression lines, and red lines are lines of equality (simulated = recovered). **B**. Recovered $\overline{\sigma_\eta^2}$ when simulated hyperparameters were set to the estimated values for the adolescent (green) and adult (purple) group, 50 times each.

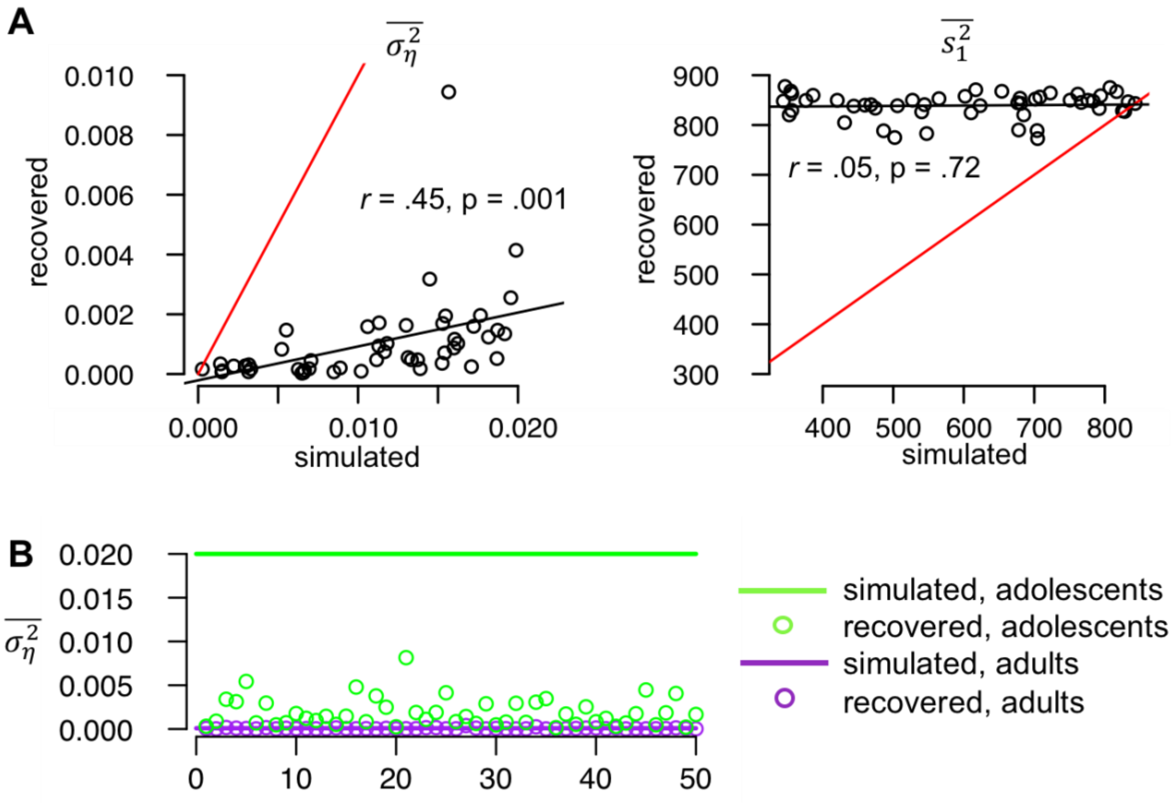

**Supplementary Fig 5**. Parameter-recovery results for the best-fitting models in the choice task. **A**. Simulated vs. recovered group-level mean parameters of the asymmetric reinforcement-learning model + dynamic softmax. **B**. Simulated vs. recovered group-level mean parameters of the reinforcement learning/Pearce-Hall hybrid model + dynamic softmax. Red lines are lines of equality.

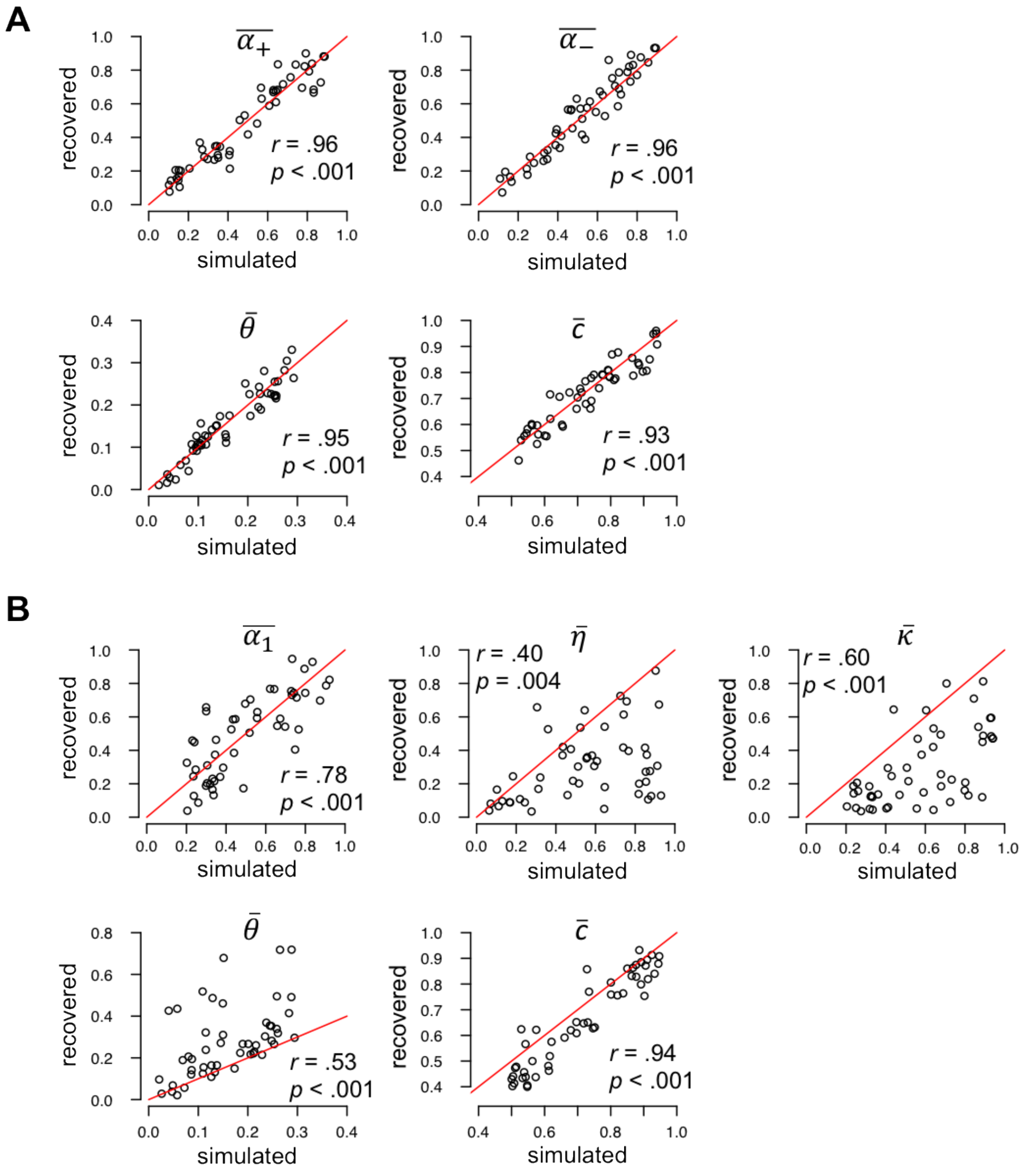

**Supplementary Table 1**. Medians (and 95% highest density intervals) of the posterior distributions shown in Fig 5A, C and D in the main text.

| | Estimation task | | | |
|---|---|---|---|---|
| | Low noise | | High noise | |
| | Adults | Adolescents | Adults | Adolescents |
| $\overline{\sigma_\eta^2}$ | .00009 (.000003 - .0004) | .02 (.0007 - .08) | .00003 (.000001 - .0001) | .001 (.00001 - .006) |
| $\overline{s_1^2}$ | 819 (469 - 1,000) | 844 (512 - 1,000) | 342 (68 – 831) | 778 (382 - 1,000) |
| | Choice task | | | |
| | Adults | | Adolescents | |
| $\overline{\alpha_1}$ | .91 (.71 – 1.0) | | | |
| $\bar{\eta}$ | .12 (.015 - .23) | | | |
| $\bar{\kappa}$ | .73 (.58 - .96) | | | |
| $\overline{\alpha_+}$ | | | .52 (.32 - .71) | |
| $\overline{\alpha_-}$ | | | .49 (.34 - .65) | |
| $\bar{\theta}$ | .22 (.18 - .27) | | .14 (.08 - .21) | |
| $\bar{c}$ | .79 (.73 - .86) | | .74 (.64 - .85) | |

**Supplementary Table 2.** Prior distributions for the simulation hyperparameters used in the model-recovery analysis.

| Model | Prior distributions |
|---|---|
| RL | $\alpha^{mean}$~U(.56, .85), $\alpha^{prec}$~ U(6, 14) |
| RL2 | $\alpha_{+}^{mean}$~U(.56, .83), $\alpha_{+}^{prec}$~ U(4, 11), $\alpha_{-}^{mean}$~U(.54, .83), $\alpha_{-}^{prec}$~ U(6, 21) |
| KF | $\sigma_{\eta}^{2\ scale}$ ~ U(.00003, .02), $s_{s,1}^{2\ scale}$~ U(342, 844) |
| PH | $\alpha_{1}^{mean}$~U(.97, .99), $\alpha_{1}^{prec}$~U(6, 29), $\eta^{mean}$~U(.25, .39), $\eta^{prec}$~U(3, 4), $\kappa^{mean}$~U(.96, .99), $\kappa^{prec}$~U(4, 33) |
| RL/con expl | $\alpha^{mean}$~U(.42, .43), $\alpha^{prec}$~ U(5, 6), $\beta^{mean}$~U(.11, .20), $\beta^{prec}$~ U(8, 6) |
| RL/dyn expl | $\alpha^{mean}$~U(.55, .59), $\alpha^{prec}$~ U(6, 8), $\theta^{mean}$~U(.12, .22), $\theta^{prec}$~ U(4, 17), $c^{mean}$~U(.72, .78), $c^{prec}$~ U(5.5, 5.8) |
| RL2/con expl | $\alpha_{+}^{mean}$~U(.25, .26), $\alpha_{+}^{prec}$~ U(7,13), $\alpha_{-}^{mean}$~U(.46, .51), $\alpha_{-}^{prec}$~ U(2.5, 2.8), $\beta^{mean}$~U(.18, .22), $\beta^{prec}$~ U(5, 45) |
| RL2/dyn expl | $\alpha_{+}^{mean}$~U(.52, .61), $\alpha_{+}^{prec}$~ U(3, 5), $\alpha_{-}^{mean}$~U(.49, .60), $\alpha_{-}^{prec}$~ U(2, 3), $\theta^{mean}$~U(.14, .22), $\theta^{prec}$~ U(3, 17), $c^{mean}$~U(.74, .79), $c^{prec}$~ U(5, 6) |
| KF/con expl | $\sigma_{\eta}^{2\ scale}$ ~ U(.04, .11), $s_{s,1}^{2\ scale}$~ U(.05, .14), $\beta^{mean}$~U(.11, .21), $\beta^{prec}$~ U(6, 52) |
| KF/dyn expl | $\sigma_{\eta}^{2\ scale}$ ~ U(.06, .28), $s_{s,1}^{2\ scale}$~ U(16, 231), $\theta^{mean}$~U(.10, .21), $\theta^{prec}$~ U(5, 16), $c^{mean}$~U(.75, .80), $c^{prec}$~ U(4.8, 5.2) |
| PH/con expl | $\alpha_{1}^{mean}$~U(.68, .71), $\alpha_{1}^{prec}$~U(4, 11), $\eta^{mean}$~U(.04, .05), $\eta^{prec}$~U(3, 7), $\kappa^{mean}$~U(.61, .62), $\kappa^{prec}$~U(4, 8), $\beta^{mean}$~U(.11, .20), $\beta^{prec}$~ U(7, 70) |
| PH/dyn expl | $\alpha_{1}^{mean}$~U(.86, .91), $\alpha_{1}^{prec}$~U(10, 16), $\eta^{mean}$~U(.07, .12), $\eta^{prec}$~U(2, 9), $\kappa^{mean}$~U(.73, .84), $\kappa^{prec}$~U(5, 9), $\theta^{mean}$~U(.11, .22), $\theta^{prec}$~ U(5, 15), $c^{mean}$~U(.73, .79), $c^{prec}$~ U(5, 6) |

Notes: The first four model were used to simulate estimation data and the last four models to simulate choice data. RL, RL2, KF, and PH are the reinforcement-learning, asymmetric reinforcement-learning, Kalman-filter, and reinforcement learning/Pearce-Hall hybrid model, respectively, dyn = dynamic, con = constant, expl = exploration, prec = precision (variance$^{-1}$)
